# Endocytosis of a zinc transceptor ZIP4 is mediated by AP2 through an atypical dileucine motif

**DOI:** 10.1101/2025.05.03.652025

**Authors:** Tianqi Wang, Chi Zhang, Yao Zhang, Jian Hu

## Abstract

Human ZIP4 is an essential zinc transporter for dietary zinc absorption and also a transceptor that undergoes zinc-dependent endocytosis to regulate cellular zinc uptake capacity in response to changes in zinc availability. However, the detailed molecular mechanism of this post-translational regulation remains elusive. In this work, we focus on the LxL motif (formerly LQL motif), which is located in the longest cytosolic loop and indispensable for ZIP4 endocytosis, to elucidate how it is involved in ZIP4 endocytosis in a zinc-responsive manner. Integrating a combination of biochemical, cell/molecular biology and modelling approaches, our data collectively support a working model that the LxL motif, which is found to be more exposed and/or less ordered upon increased zinc availability, acts as an atypical dileucine motif to interact with the adaptor protein complex 2 for clathrin-mediated endocytosis. These findings provide insights into the molecular basis of zinc-dependent ZIP4 endocytosis, advancing the understanding of cellular zinc homeostasis and providing a paradigm for mechanistic studies of substrate-induced endocytosis of other nutrient transceptors.

**Highlights:**

- Hydrophobicity of the LxL motif is essential for ZIP4 endocytosis
- Endocytosis of endogenously expressed ZIP4 depends on Clathrin and AP2
- The LxL motif binds to the σ2 subunit of AP2
- Cellular zinc levels regulate the structure of the LxL motif

## INTRODUCTION

Some nutrient transporters act as transceptors, integrating substrate transport and substrate sensing to better maintain cellular nutrient homeostasis.^1-3^ For many, but not all, transceptors, the substrate sensing function is manifested by substrate-induced endocytosis, in which a transceptor is removed from the plasma membrane when the level of substrate or substrate surrogate is elevated,^2,4-18^ allowing cells to rapidly adjust transport capacity in response to changes in nutrient availability. Unravelling the molecular basis of this post-translational regulatory mechanism for transceptors is crucial for understanding nutrient homeostasis under physiological and pathological conditions.^19,20^

Among the known transceptors, ZIP4, a representative Zrt-/Irt-like protein (ZIP) metal transporter that is specifically expressed in the small intestine and the kidney for zinc uptake from food and zinc reabsorption from urine,^20-25^ is a central player in maintaining systemic zinc homeostasis. At the cellular level, the zinc-sensing function of ZIP4 enables dynamic regulation of its cell surface expression, ensuring sufficient, but not excessive, zinc uptake for critical biological processes. ZIP4 undergoes internalization through two distinct mechanisms in response to changes in zinc availability. When zinc concentration rises significantly above physiological levels (reaching tens to hundreds of micromolar), the second intracellular loop (hereafter the IL2 loop) of ZIP4 is ubiquitinated for subsequent degradation in both lysosome and proteasome.^26^ Zinc sensing in this process is conducted by the histidine-rich segment located in the IL2 loop.^27^ This long and highly variable loop has also been shown to be important for the non-iron metal-induced endocytic degradation of a plant iron-transporting ZIP, IRT1.^4,28^ In contrast, at physiological zinc concentrations (low micromolar range), ZIP4 undergoes constitutive, ubiquitination-independent endocytosis (referred to as zinc-dependent endocytosis).^29^ Upon zinc depletion, this constitutive endocytosis is significantly reduced, whereas transferrin internalization is unchanged under the same condition, supporting that the observed changes in endocytosis is specific for ZIP4.^30^ By scanning potential zinc-binding sites, we showed that the transport site of human ZIP4 functions as a zinc-sensing site.^30^ An extracellular zinc binding site at the dimerization interface of mouse ZIP4 has also been suggested to play a role in zinc sensing.^31^ While it has been proposed that zinc binding to ZIP4 leads to conformational changes that allow ZIP4 to engage the endocytic machinery for internalization, the exact structural changes remain to be elucidated.

During membrane protein endocytosis, cargo recognition by adaptor protein is a critical step that involves specific protein-protein interactions. Various sorting signals have been identified for different endocytic mechanisms.^32^ In clathrin-mediated endocytosis, the heterotetramer adaptor protein complex 2 (AP2, composed of α, β2, µ2, and σ2 subunits) plays a central role in cargo selection by either direct binding to cargo proteins via sorting motifs or by binding to cargo-specific adaptor proteins that bridge AP2 to cargo proteins.^33-35^ The best-characterized sorting motifs that are recruited by AP2 include the tyrosine-based motif (YxxØ, where Ø is a bulky hydrophobic amino acid, and [F/Y]xNPx[Y/F]) and the dileucine motif ([D/E]xxxL[L/I]),^32,34^ while many other sorting signals have been identified and more are expected to exist.^34,36^ In our previous study on human ZIP4 endocytosis, we systematically scanned the IL2 loop and identified a conserved ^452^LQL^454^ segment essential for ZIP4 endocytosis.^30^ However, the precise role of this segment in zinc-dependent ZIP4 endocytosis remains unclear.

In this study, we employed multidisciplinary approaches to investigate the role and structural changes of the LQL segment in ZIP4 endocytosis. Using a flow cytometry-based antibody uptake assay, we demonstrated that the hydrophobicity of the two leucine residues is essential for ZIP4 internalization. Using a custom anti-ZIP4 monoclonal antibody (mAb), we found that endogenously expressed ZIP4 in HepG2 cells undergoes a clathrin-mediated and AP2-dependent endocytosis. Importantly, our data support a model in which the LxL motif binds to the hydrophobic pocket of the σ2 subunit of AP2 – a site typically used for the binding of canonical dileucine motifs, suggesting that it functions as an atypical dileucine sorting signal in cargo recognition. Cysteine accessibility assays performed on a series of single cysteine variants revealed that the IL2 loop undergoes significant zinc-dependent conformational changes, providing a structural basis for zinc sensing. Together, these findings provide mechanistic insights into the molecular basis of ZIP4 endocytosis.

## RESULTS

### Essential roles of the leucine residues in the LQL segment

Our previous study has shown that alanine substitution of the residues in the LQL segment diminished ZIP4 endocytosis.^30^ In this work, we conducted an extensive mutagenesis study on these residues to clarify what properties of the amino acids at these positions are important. To do so, the C-terminal HA-tagged human ZIP4 and its variants were transiently expressed in HEK293T cells (**Figure 1A**). After incubation with the anti-HA antibody at 37°C to allow antibody uptake, cell lysates were applied to Western blot to detect the internalized antibody (**Figure 1B)**. For the first leucine (L452), replacement with either isoleucine or methionine did not affect antibody uptake, but alanine or asparagine substitution nearly abolished antibody uptake. Similarly, alanine or asparagine substitution of the second leucine (L454) led to largely reduced antibody uptake. Although the substitution with methionine did not affect endocytosis, the substitution by isoleucine significantly reduced it. In contrast, none of the substitutions on the glutamine residue (Q453) reduced antibody uptake. We also tested substitutions on E457, a generally conserved residue close to the LQL segment, and the results indicated that amino acid replacement at this position has no effect on ZIP4 endocytosis.

**Figure 1.**
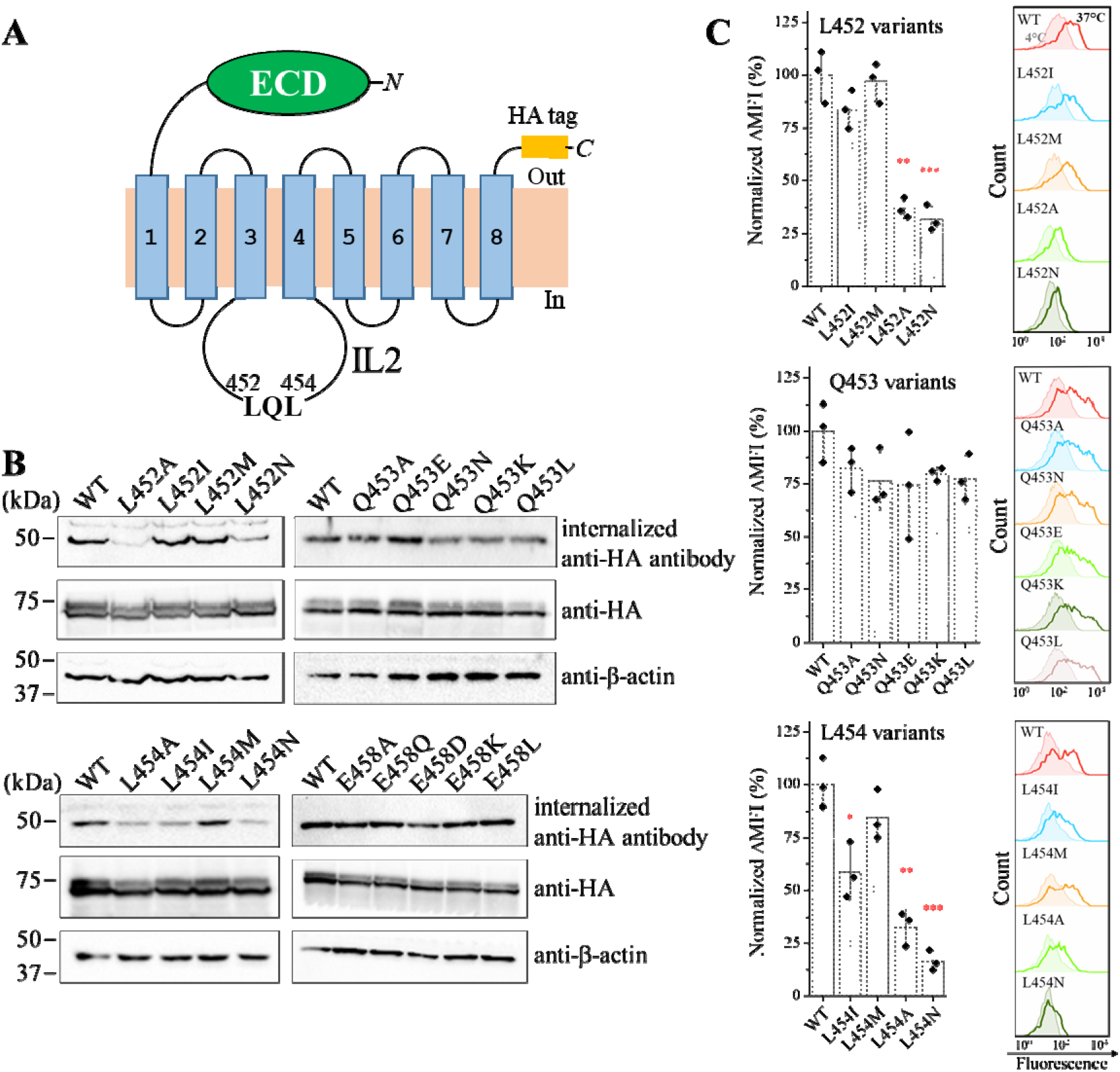
Effects of the LQL mutations on human ZIP4 endocytosis in HEK293T cells. (**A**) Topology of hZIP4-HA in the plasma membrane. The LQL segment is located in the IL2 loop. The ECD is shown as a green oval. The HA tag (orange rectangle) is exposed to the extracellular space. (**B**) Effects of substitutions of the residues in the LQL segment on human ZIP4-HA endocytosis. E458 is a generally conserved residue in ZIP4 and thus was tested along with the LQL sequence. The internalized mouse anti-HA antibodies were detected by Western blot using an HRP-conjugated anti-mouse antibody. The total expression levels of ZIP4 variants were detected by an anti-HA antibody. Western blot of anti-β actin was used as loading control. (**C**) Comparison of anti-HA antibody uptake between wild-type ZIP4-HA and its variants using a flow cytometry-based assay. ΔMFI was calculated by subtracting the MFI of the cells incubated with the antibody at 4°C from the MFI of the cells incubated with the antibody at 37°C. The values of ΔMFI were then normalized by setting the value for wild-type ZIP4-HA as 100%. Each column chart shows the data in one of three independent experiments with three biological replicates tested for each condition. The representative histograms on the right show the profiles of 4°C (shaded) and 37°C (open). Statistical analyses were performed using two-tailed Student’s *t*-test. The *P* values are 0.0011 (L452A) 0.00099 (L452N), 0.015 (L454I), 0.0012 (L454A), and 0.00033 (L454N), respectively. *: *P*<0.05; **: *P*<0.01; ***: *P*<0.001.

Given that the Western blot-based assay barely allows a statistical analysis, we applied a flow cytometry-based antibody uptake assay to study ZIP4 endocytosis (**Figure S1**). In this assay, cells transiently expressing ZIP4 or its variants were incubated with a fluorescence-labelled anti-HA antibody at 4°C or 37°C for 45 min and then washed with an acidic buffer (pH 4.0) at 4°C to remove the cell surface bound antibody before flow cytometry analysis. The differences in mean fluorescence intensity (MFI) between the samples treated at 4°C and 37°C represent the signals derived from the internalized antibodies. Consistent with the results obtained from the Western blot-based assay, antibody uptake was significantly reduced for the L452A, L452N, L454A, and L454N variants (**Figure 1C**), indicating that the hydrophobicity of L452 and L454 is important for ZIP4 endocytosis. Substitution of L454 with isoleucine significantly reduced endocytosis, whereas substitution with methionine had little effect, suggesting that a terminal methyl group on the side chain is critical. As the results of the Q453 variants showed that changes in charge, size and polarity at this position did not affect ZIP4 endocytosis, the LQL segment is henceforth be referred to as the LxL motif.

### Development of an anti-ZIP4 mAb to recognize endogenously expressed ZIP4

Chlorpromazine, an endocytosis inhibitor believed to disrupt the formation of clathrin-coated pits,^37^ was shown to dose-dependently reduce ZIP4 endocytosis in HEK293T cells,^30^ suggesting clathrin-mediated endocytosis. Given that the endocytosis mechanism for overexpressed proteins may differ from that for endogenously expressed proteins, we developed an anti-human ZIP4 mAb (Clone 1G3B2) to study ZIP4 endocytosis in HepG2 cells, a human hepatocyte cancer cell line in which ZIP4 mRNA and protein expression have been documented in Human Protein Atlas^38^ and ProteomicsDB,^39^ respectively. The mouse-derived 1G3B2 mAb, generated using the purified extracellular domain (ECD) of human ZIP4 as antigen (**Figure S2A**), stained the live HEK293T cells overexpressing ZIP4, whereas the empty vector-transfected cells showed no staining in immunofluorescence imaging (**Figure S2B**), indicating that the 1G3B2 mAb specifically recognizes human ZIP4 in its native state.

### Clathrin-mediated and AP2-dependent ZIP4 endocytosis

We then used the 1G3B2 mAb to stain the live HepG2 cells and the result confirmed the expression of ZIP4 in this cell line and also showed that the antibody was internalized when cells were incubated at 37°C, but barely at 4°C (**Figure 2A**). Using the similar flow cytometry-based antibody uptake assay, the MFI difference between the cells incubated at 37°C and 4°C was found to increase over time (**Figure S2C**). To confirm the antibody uptake is ZIP4-specific, ZIP4 was knocked down by RNA interference and the greatly diminished antibody uptake indicated that the 1G3B2 mAb can be used as a tracer to study ZIP4 endocytosis in HepG2 cells (**Figure 2B**).

**Figure 2.**
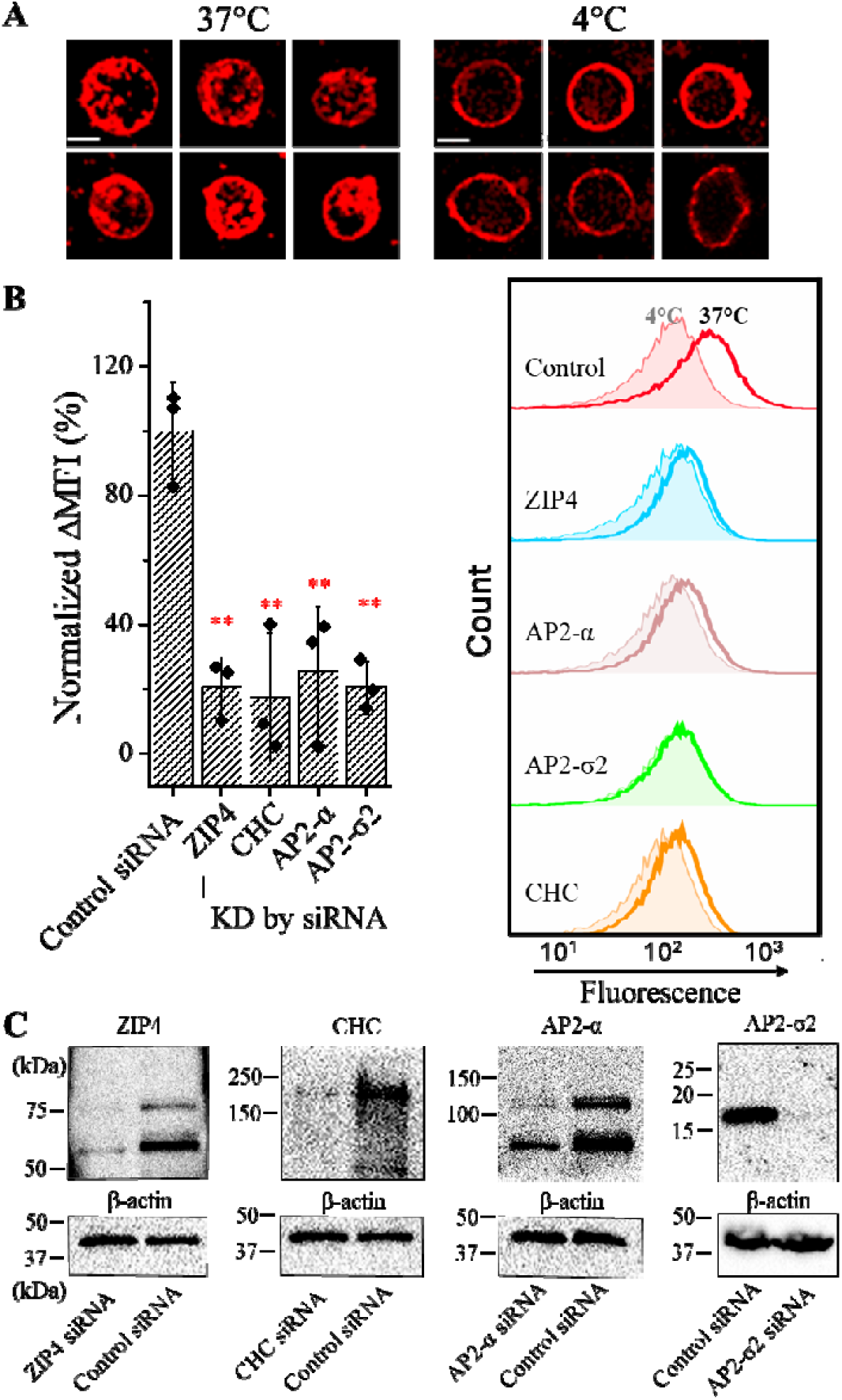
Endocytosis of endogenously expressed ZIP4 in HepG2 cells. (**A**) Internalization of the 1G3B2 anti-ZIP4 mAb by HepG2 cells. Live HepG2 cells were incubated with 1G3B2 mAb at 4°C and 37°C, respectively. After wash, fixation and permeabilization, cells were stained with an Alexa Fluor-568 conjugated anti-mouse antibody. Immunofluorescence images were taken using a confocal microscopy. Representative cells from multiple images were shown. Scale bars indicate 10 μm. (**B**) Analysis of 1G3B2 mAb uptake by flow cytometry. The target proteins were knocked down by siRNA. ΔMFI was calculated by subtracting the MFI of the cells incubated with the antibody at 4°C from the MFI of the cells incubated with the antibody at 37°C. The values of ΔMFI were then normalized by setting the value of the control group as 100%. The column chart shows the data in one of two independent experiments with three biological replicates tested for each condition. The representative results are shown in histogram chart on the right. Statistical analyses were conducted using two-tailed Student’s *t*-test. The *P* values are 0.0068 (ZIP4), 0.0013 (CHC), 0.0047 (AP2-α), and 0.0017 (AP2-σ2), respectively. **: *P*<0.01. (**C**) Knockdown of the target proteins by control siRNA or specific siRNA. The expression levels were analyzed in Western blot using the corresponding antibodies.

To test the possible involvement of clathrin in ZIP4 endocytosis in HepG2 cells, as suggested by the inhibition of ZIP4 endocytosis in HEK293T cells by chlorpromazine,^30^ we knocked down clathrin heavy chain using siRNA and found the 1G3B2 mAb uptake was largely abolished (**Figure 2B**), indicating that ZIP4 endocytosis is a clathrin-mediated process. Since AP2 is a central player in clathrin-mediated endocytosis, we tested whether it is required for ZIP4 endocytosis by knocking down the α and σ subunits using siRNAs. The flow cytometry data showed drastically reduced antibody uptake (**Figure 2B**), indicating that AP2 is essential for ZIP4 endocytosis. Given that AP2 may directly bind to the cargo protein or act indirectly by binding to a cargo-specific adaptor protein that recognizes the cargo protein,^34^ these results raised a question of the exact role of AP2 in ZIP4 endocytosis.

### Predicted structural model of the AP2-IL2 complex

To look for a potential binding between ZIP4 and AP2, we generated structural models of the AP2-IL2 complex using AlphaFold 3, where a 57-aa peptide corresponding to the IL2 loop of human ZIP4 (residue 438-494) was used (**Figure 3A**). AP2 in the AlphaFold generated structure models adopt a consistent open-like conformation when compared to the experimentally solved open state AP2 complex structures (**Figure S3**).^40-42^ In 3 of 35 generated models, the sequence of ^476^ESPELL^482^ was found to bind to a hydrophobic pocket in the σ2 subunit that is known for binding of the canonical dileucine motif.^43^ However, our previous mutagenesis study has shown that alanine substitutions of the leucine residues in this segment had no effect on ZIP4 endocytosis.^30^ Of great interest, 14 models consistently showed the binding of the LxL motif to the same hydrophobic pocket in the σ2 subunit (**Figure 3A**). Structural comparison revealed a similar mode for the binding of the canonical dileucine motif (a CD4-derived peptide, PDB IDs: 2JKR and 6QH6)^43,44^ and the LxL motif to the σ2 subunit (**Figure 3B**) – the first leucine residue (L8 in the CD4 peptide and L452 in ZIP4) occupies the same position deep in the pocket; the second leucine residue (L9 in CD4 and L454 in ZIP4) binds to a shallow position at the entrance of the pocket and the isopropyl groups on their side chains are superimposable. Structural differences are also remarkable. The two consecutive leucine residues in CD4 form a sharp turn in order to position their side chains on the same side of the backbone. In contrast, due to the x residue inserted between the two leucine residues, the backbone of the LxL motif adopts an extended conformation, positioning the side chains of the two leucine residues on the same side of the backbone for the binding to the hydrophobic pocket. As a result, the x residue faces away from the binding pocket and thus is not involved in the binding to the σ2 subunit. Overall, the AlphaFold3 predicted structures raised a possibility that the LxL motif may bind to the hydrophobic pocket of the σ2 subunit.

**Figure 3.**
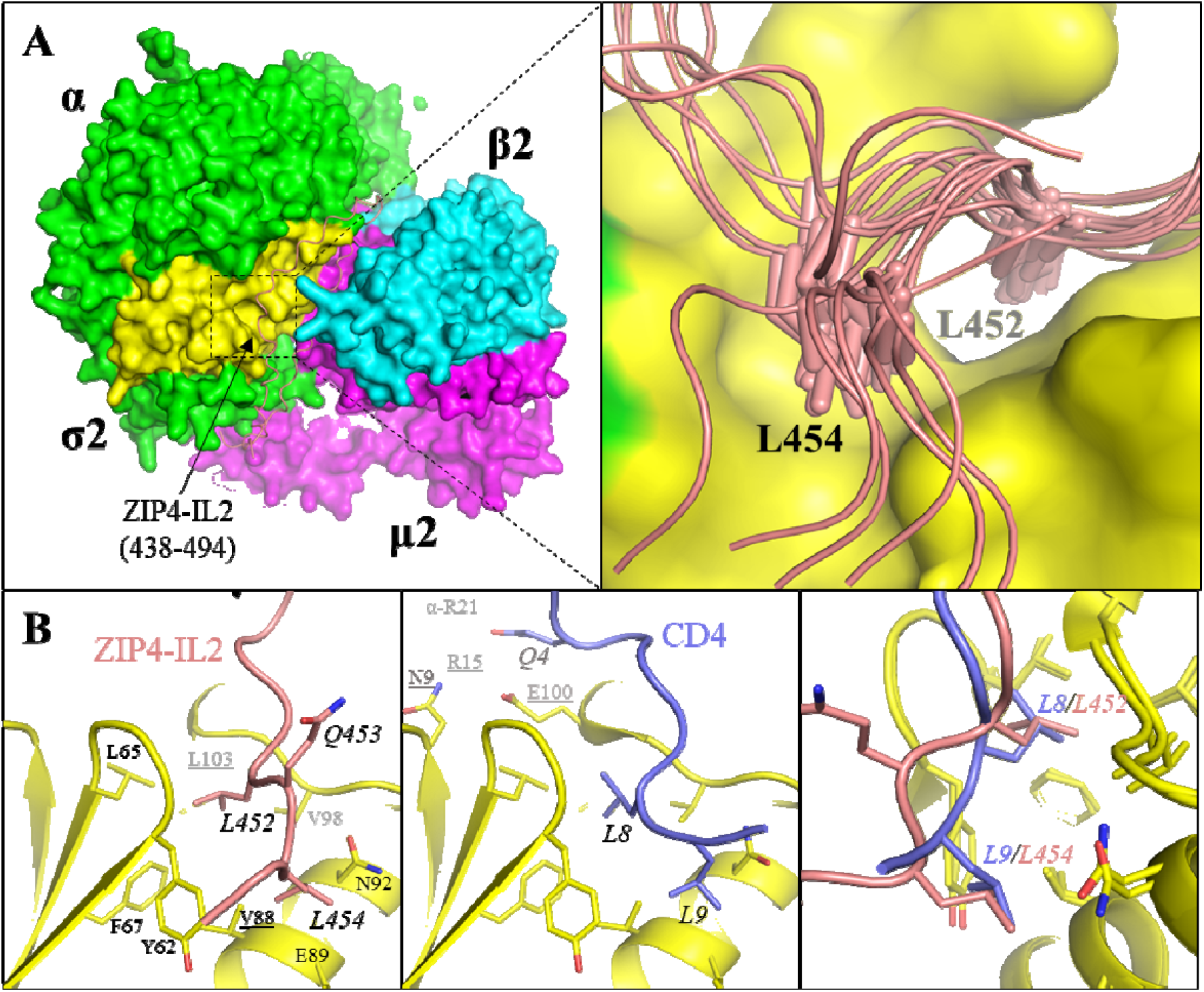
Structural model of the AP2-IL2 complex generated by AlphaFold 3. (**A**) Overall structure (*left*) and zoomed-in view (*right*). AP2 is shown in surface mode and the ZIP4-IL2 peptide (438-494, pink) is in cartoon mode (*left*). 14 of 35 predicted models showed a consistent binding of the LxL motif to a hydrophobic pocket in the σ2 subunit (*right*). L452 and L454 are depicted in stick mode. (**B**) Comparison of the binding of the LxL motif (*left*) and the canonical dileucine motif in CD4 (*middle*, PDB: 2JKR). The superimposed structures are shown in the right panel. The residues mutated in this work (V88, L103, N9, R15, and E100 in the σ2 subunit) are underlined.

### Essential role of the dileucine motif-binding pocket in ZIP4 endocytosis

To confirm the importance of the σ2 subunit in ZIP4 endocytosis, HepG2 cells were co-transfected with the siRNAs, which knock down the endogenously expressed σ2 subunit, and a vector harboring the gene encoding an siRNA-resistant wild-type σ2 protein. The results showed that overexpression of the wild-type σ2 protein not only rescued the attenuated ZIP4 endocytosis, but also increased the endocytosis level even higher than that in the cells transfected with control siRNA (**Figure 4A**). Since the hydrophobicity of the pocket in the σ2 subunit has been shown to be important for the binding of canonical dileucine motif, we applied two functionally compromised variants (V88D and L103S) to the rescue experiment.^43^ Compared to the cells overexpressing the wild-type σ2 protein, the cells expressing either of these variants showed a significantly reduced rescue effect, and when the two mutations were combined (V88D/L103S), the rescue effect was totally eliminated (**Figure 4A**). These results strongly indicated an essential role of the dileucine motif-binding pocket in ZIP4 endocytosis.

**Figure 4.**
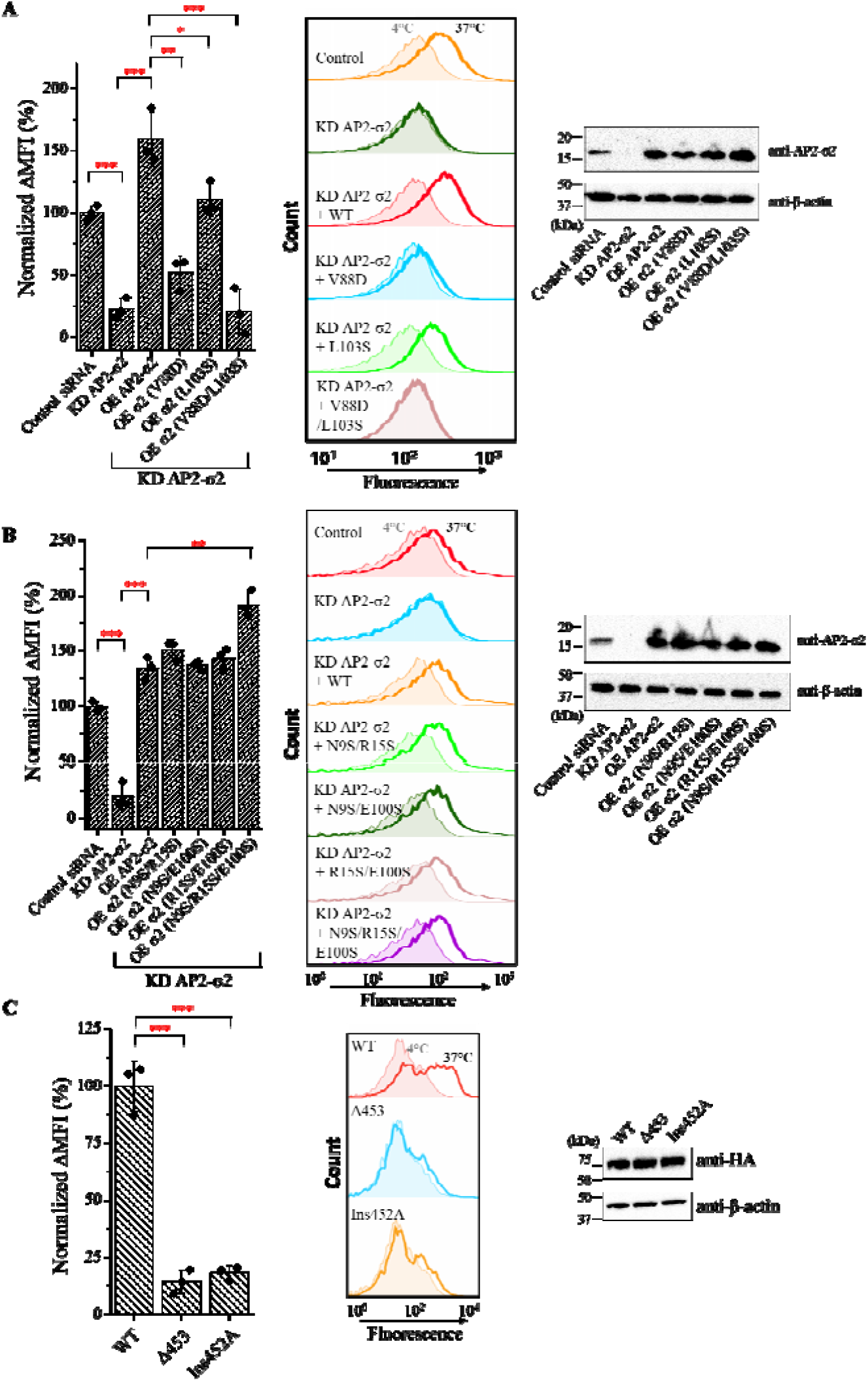
Identification of the key residues in the AP2-IL2 interaction. (**A**) The importance of the hydrophobic pocket in the σ2 subunit. The σ2 subunit in HepG2 cells was knocked down by siRNA and rescued by overexpressing an siRNA-resistant σ2 subunit or its variants with hydrophobic residues (V88 and L103) substituted with polar/charged amino acids. ΔMFI was calculated by subtracting the MFI of the cells incubated with the antibody at 4°C from the MFI of the cells incubated with the antibody at 37°C. The values of ΔMFI were then normalized by setting the value of the control group as 100%. The representative results are shown in histogram chart (*middle*). Statistical analyses were conducted using two-tailed Student’s *t*-test. The *P* values from left to right are 0.00020, 0.00056, 0.0020, 0.031, and 0.0011, respectively. *: *P*<0.05; **: *P*<0.01; ***: *P*<0.001. The expression levels of the wild-type σ2 subunit or its variants were examined by Western blot, and β-actin was used as loading control. (**B**) Examination of the involvement of the electrostatic patch (composed of N9, R15, and E100) in the σ2 subunit. The knockdown and rescue experiments and Western blot were conducted in the same as in (A). The *P* values from left to right are 0.00043, 0.00027, and 0.0035, respectively. (**C**) Endocytosis of wild-type ZIP4-HA and its variants with changed distance between the two leucine residues in the LxL motif. The constructs were transiently expressed in HEK293T cells and the anti-HA antibody uptake assay was conducted and detected as described in Figure 1C. For **A, B**, and **C**, the column chart (*left*) shows the data in one of two independent experiments with three biological replicates tested for each condition.

Comparing the binding modes of the canonical dileucine motif and the LxL motif to the σ2 subunit (**Figure 3B**), one significant difference is that the latter does not involve an acidic residue that binds to a positively charged patch of the σ2 subunit through electrostatic interactions. To experimentally examine the role of this charged surface in ZIP4 endocytosis, we tested additional σ2 variants in which the residues that form the electrostatic interactions with the acidic residue (or a glutamine in the case of CD4) in the canonical dileucine motif (N9, R15, and E100) were replaced with serine residues (**Figure 4B**). Previous studies have shown that disruption of this interface reduces the binding affinity for CD4-derived peptide,^43^ but rescue experiments using these variants showed that none of them compromised the rescue efficacy, further supporting that the binding of the LxL motif to the hydrophobic pocket of the σ2 subunit does not need additional electrostatic interactions to stabilize it. This result also does not support the possibility that AP2 may bind to a yet identified ZIP4-specific adaptor that uses a canonical dileucine motif for AP2 engagement.

If the LxL motif does indeed bind to the hydrophobic pocket of the σ2 subunit (**Figure 3**), we wondered whether a change in the distance between the two leucine residues could be tolerated. To test this, we generated two ZIP4-HA variants, Δ 453, in which Q453 was deleted, and Ins452A, in which an alanine was inserted after L452, and expressed them in HEK293T cells for a flow cytometry-based antibody uptake assay. As shown in **Figure 4C**, neither variant was efficiently internalized, indicating that an LL sequence (in the Δ453 variant) or an LAQL sequence (in the Ins452A variant) is unable to support ZIP4 endocytosis, likely due to impaired binding to the σ2 subunit.

Taken together, the rescue and mutagenesis experiments consistently supported the model that the σ2 subunit of AP2 binds to the LxL motif in the IL2 loop of ZIP4. We then looked at how this interaction is regulated in a zinc-dependent manner.

### Zinc-dependent conformational changes of the IL2 loop

As reported in our early study, adding zinc to purified ZIP4 increased the resistance of the IL2 loop to chymotrypsin digestion in a dose-dependent manner.^30^ To study the zinc binding-induced conformational changes of ZIP4, we conducted a cysteine accessibility assay on mouse ZIP4 expressed in live HEK293T cells. Mouse ZIP4 was chosen in this experiment is because it appears to be more stable than human ZIP4 in HEK293T cells according to the previous mutagenesis studies and therefore is thought to be more tolerant to multiple mutations.^20,45^ As there are 15 endogenous cysteine residues in the wild-type mouse ZIP4, including eight in the ECD forming four disulfide bonds as shown in the structure of ZIP4-ECD^46^ and seven in the transmembrane domain (TMD), we removed all of them by deleting the ECD, which is not required for ZIP4 endocytosis or zinc sensing,^30,31^ and replacing the seven cysteine residues in the TMD with alanine, generating a variant named cysteine-free TMD (hereafter CysFree-TMD). Cell-based transport assay indicated that CysFree-TMD is a functional Zn^2+^ transporter and undergoes endocytosis as wild-type ZIP4 (**Figure S4**). Then, cysteine residues were introduced to generally evenly distributed positions in the IL2 loop of CysFree-TMD with no substitutions on conserved residues, resulting in eight variants with one cysteine in each of them (**Figure 5A**).

**Figure 5.**
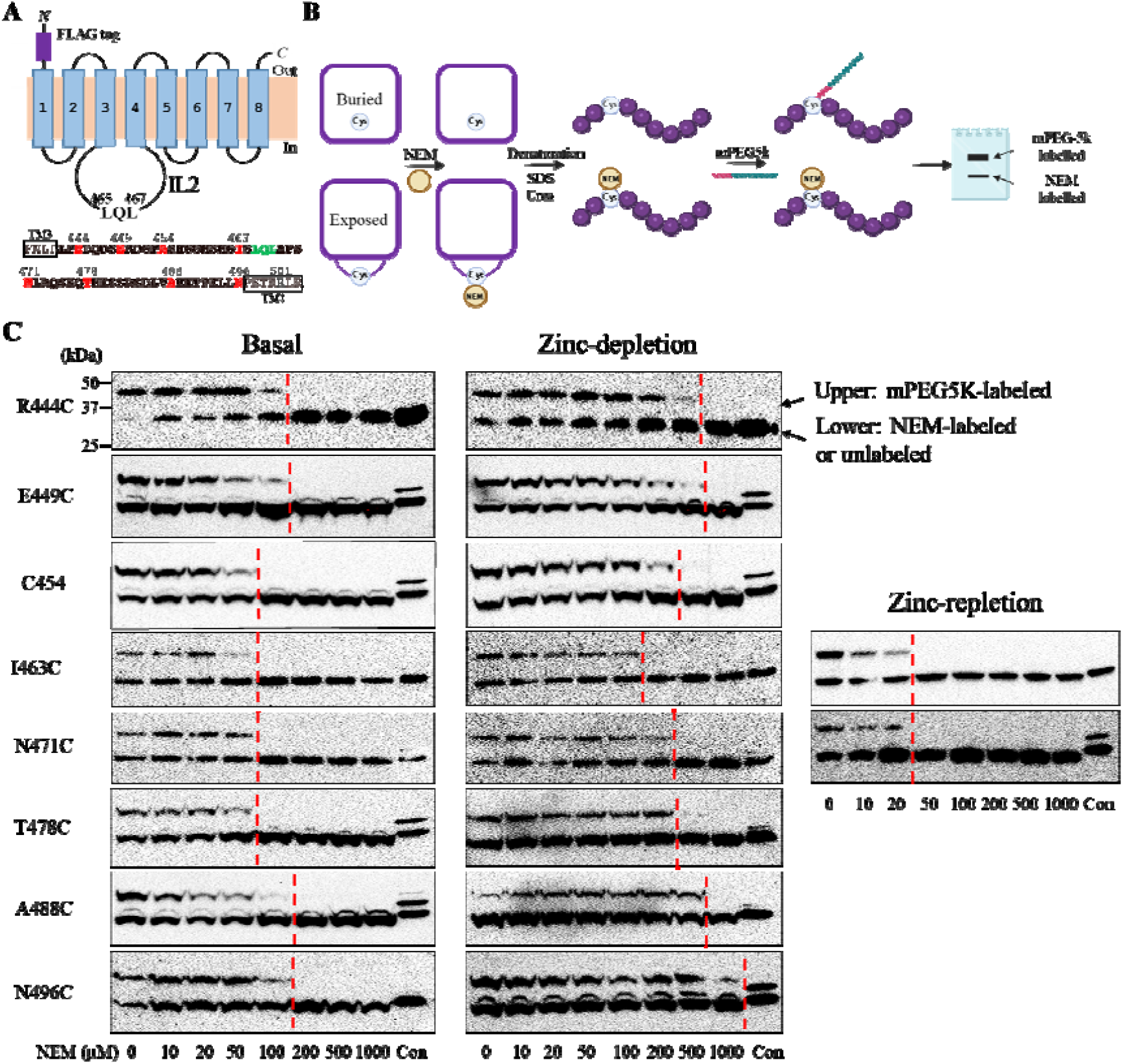
Cysteine accessibility assay of mouse ZIP4 expressed in HEK293T cells. (**A**) Topology of the cysteine-free TMD of mouse ZIP4 (CysFree-TMD). The amino acid sequence of the IL2 loop is bolded and the residues substituted with cysteines are highlighted in red. The LxL motif is shown in green. Note that a cysteine residue is present at the position of 454 in wild-type ZIP4. (**B**) Cartoon illustration of the experimental procedure. (**C**) Cysteine accessibility assays of eight single cysteine variants. For each variant, an NEM titration was conducted and the red dashed line indicates the concentration of NEM above which the cysteine of interest is 100% labelled by NEM. Cells in control group (Con) were not treated with NEM or mPEG5k. An anti-FLAG antibody was used in Western blot experiments. The shown data are the results of one of two independent experiments with similar results.

The variants were transiently expressed in HEK293T cells and the accessibility of the introduced cysteine was studied on live cells under basal, zinc depletion, and zinc repletion conditions (**Figure 5B**). Cells expressing N-terminal FLAG-tagged CysFree-TMD variants were first incubated with a membrane-permeable thiol-reacting reagent, N-ethylmaleimide (NEM), to allow the exposed cysteine residues to be labeled by NEM. After removal of residual NEM by extensive washing, cells were lysed in a denaturation/reducing solution to completely unfold proteins to expose buried cysteine residues for the subsequent reaction with excessive monofunctional PEG-Maleimide 5K (mPEG5K), a thiol-reacting reagent with a molecular weight of 5 kDa. The resulting mixtures were then analyzed in Western blot to differentiate the exposed cysteine residues, which showed no band shift due to the low molecular weight of NEM, from the buried cysteine residues, which showed a 5 kDa increase in molecular weight due to the PEG conjugation. The ratio of the two bands reflects the extent to which the cysteine residue of interest is exposed. NEM titration was conducted for each variant for better comparison.

As shown in **Figure 5C**, the combined treatment of TPEN, a membrane-permeable high-affinity Zn^2+^ chelator, and the Chelex-treated cell culture medium (zinc depletion condition) consistently increased the mPEG5k-labeled species for all tested variants when compared to the cells maintained in the regular culture medium that contains approximately 5 µM zinc (basal condition), indicating that zinc depletion reduced the reaction of cysteine residues with NEM. To test the effects of zinc repletion, two variants, I463C and N471C, were chosen due to their close proximity to the ^465^LxL^467^ motif in mouse ZIP4. To increase cellular zinc levels, cells were first incubated with 10 µM ZnCl_2_ and 10 µM pyrithione, a zinc ionophore,^47^ and then subjected to cysteine accessibility assay. Consistent with the trend revealed under the zinc depletion conditions, zinc supplementation reduced the mPEG5k-labeled species for both variants. Therefore, this set of experiments showed that there is an inverse correlation between zinc availability and cysteine labeling by mPEG5k. Since the latter takes place only when a cysteine residue fails to react with NEM under native condition, this result indicates that the increased cellular zinc levels promote the reaction of cysteine residues with NEM. It is known that zinc ion can form a complex with thiolate and reduce the reactivity of the coordinated cysteine residue,^48^ and therefore, an increased zinc level would reduce NEM labeling. However, an opposite trend was observed, and it can be explained only when the IL2 loop underwent a zinc-responsive conformational change: it is more ordered and/or less exposed under zinc depletion conditions, whereas under zinc repletion conditions, it becomes less ordered and/or more exposed. In addition, the different extents of accessibility for the cysteine residues along the IL2 loop suggested that the LxL motif is located in a region that is least ordered and/or most exposed in the IL2 loop, supporting the role of this motif in the engagement with binding partners, such as the endocytic machinery. Overall, the results of cysteine accessibility assays provided insights into the structural response of the IL2 loop to the changes in cellular zinc levels.

## DISCUSSION

Substrate-induced endocytosis of a transceptor provides a rapid regulatory mechanism for maintaining nutrient homeostasis. How the transporter engages the endocytic machinery in a substrate-responsive manner is a central question in the study of transceptors, and various mechanisms have been proposed for different systems.^4,5,7,10,11,13,14,17^ As a zinc transceptor, ZIP4 undergoes constitutive endocytosis in a zinc-dependent manner,^29,30^ but the molecular basis of this multi-step event remains to be elucidated. In this work, by integrating cell biology, biochemistry and modelling approaches, along with a custom anti-ZIP4 mAb for the study of endogenously expressed ZIP4 in HepG2 cells, we found that the LxL motif located in the IL2 loop of ZIP4 acts as a sorting signal that adopts different conformational states in response to cellular zinc levels and interacts with AP2 for clathrin-mediated endocytosis.

Several lines of evidence supported a key role of the LxL motif in ZIP4 recognition by the endocytic machinery (**Figure 6**). First, the mutagenesis study indicated that the hydrophobicity of the two leucine residues is required for ZIP4 endocytosis, whereas the residue in between is dispensable (**Figure 1**). Second, the knockdown experiments indicated that endocytosis of endogenously expressed ZIP4 in HepG2 cells requires both clathrin and AP2 (**Figure 2**). Third, despite the insertion of a residue between the two leucine residues and the presence of a dileucine motif-like segment (^476^ESPELL^482^) in the IL2 loop, AlphaFold 3 preferentially modelled the binding of the LxL motif to the hydrophobic pocket in the AP2-σ2 subunit, which is known for the binding of the canonical dileucine motif [(D/E)xxxL(L/I)] (**Figure 3**). Fourth, the key role of this dileucine motif-binding pocket was demonstrated in the knockdown-rescue experiments (**Figures 4A**). Fifth, disruption of a polar surface used for electrostatic interactions with the acidic residue in the canonical dileucine motif did not affect ZIP4 endocytosis (**Figure 4B**). And lastly, changing the distance between the two leucine residues was not tolerated, which is consistent with the predicted binding mode where the first leucine residue occupies a deep position while the second one can only bind at the shallow position due to the small size of the binding pocket (**Figure 4C**). These data collectively suggest that the dileucine motif binding pocket in the σ2 subunit can be used to bind other small hydrophobic motifs, including the LxL motif in ZIP4. A tangentially related example is the interaction between Nef, an human immunodeficiency virus (HIV) protein that bridges target proteins to the AP1/2 complexes to disrupt protein trafficking in T cells,^49^ and SERINC5, a human protein exerting an anti-HIV activity. It has been reported that L350/I352 in SERINC5 is critically involved in the interaction with Nef,^50,51^ and this LxI sequence has been proposed to bind to the same hydrophobic pocket used to accommodate the dileucine motif in CD4, a primary target protein of Nef.^52^ Nevertheless, to further establish the binding mode of the LxL motif to AP2, a structural study is warranted for the future research. Another issue to be addressed is the sufficiency of the LxL motif for ZIP4 to be recruited by AP2. Although this motif is essential for ZIP4 endocytosis, our previous study showed that the LQL sequence fused to the cytoplasmic tail of CD8 did not induce internalization of the chimeric protein.^30^ We therefore speculate that in addition to the LxL motif, other structural elements of ZIP4 are critical for the engagement of ZIP4 with AP2. Alternatively, the LxL motif alone is sufficient to be recruited by AP2 only when it is within an appropriate distance from the TMD of ZIP4.^34,53^

**Figure 6.**
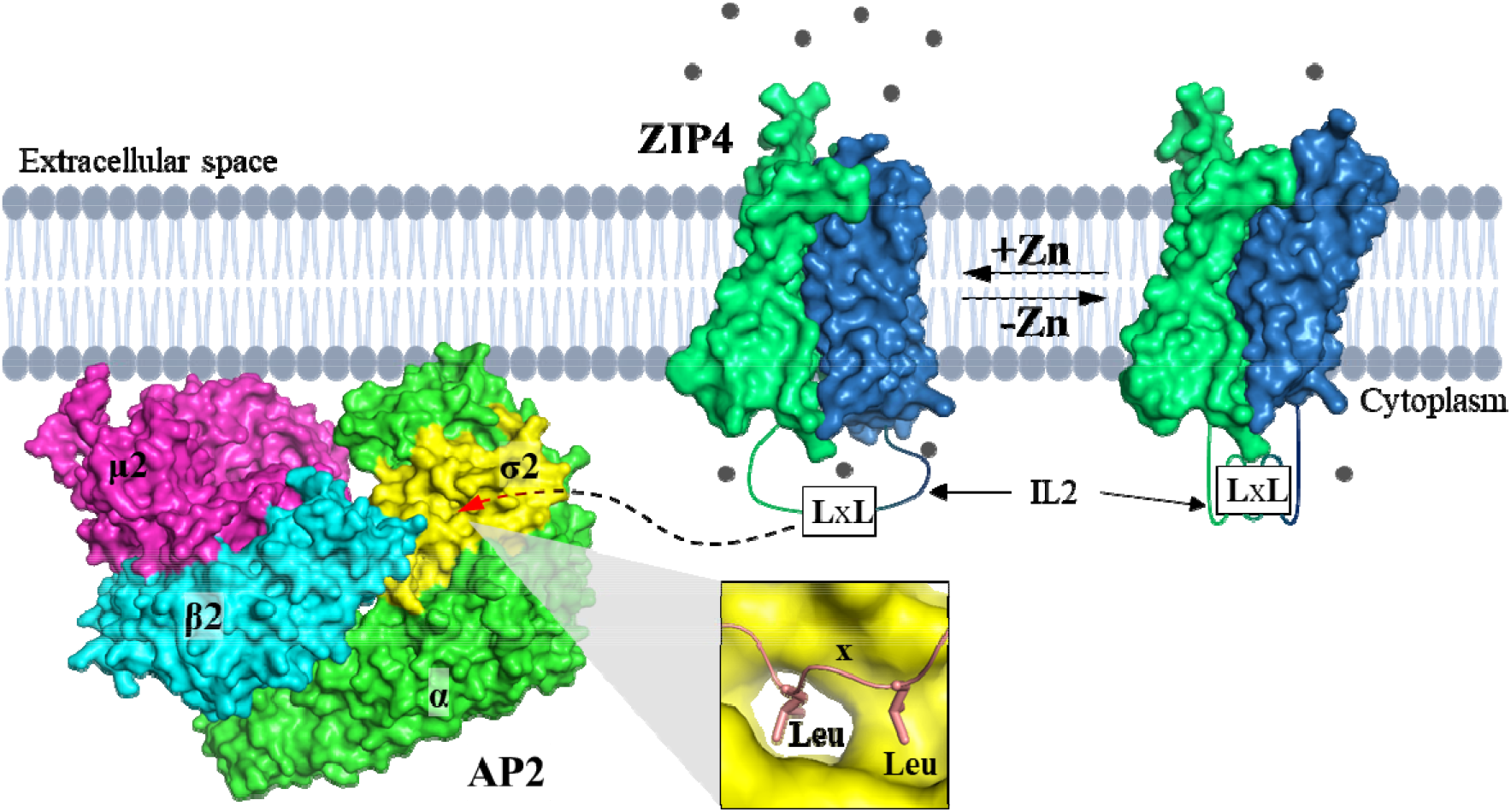
Proposed mechanism of zinc-dependent ZIP4 endocytosis. Cellular zinc availability regulates the conformational states of the IL2 loop, where the LxL motif is located. Under zinc depletion conditions, the LxL motif is less accessible (right); with increased zinc levels, the LxL motif becomes more exposed (left), allowing it to be recruited by AP2 (PDB: 2XA7, orientated in the membrane-bound state) through a direct binding to the hydrophobic pocket in the σ2 subunit (indicated by arrow and zoomed in the inset), where canonical dileucine motifs of cargo proteins bind. For clarity, the ECD of ZIP4 is trimmed. The transport domain of ZIP4 is depicted as a two-domain protein with the scaffold and transport domains shown in green and blue, respectively. Alternating access is achieved by vertical sliding of the transport domain against the static scaffold domain, which is proposed to trigger conformational changes of the IL2 loop. Zinc ions are depicted as grey spheres.

Consistent with our previous study on purified ZIP4, the systematic cysteine accessibility assays conducted on multiple positions in the IL2 loop indicated that the IL2 loop adopts distinct conformations in response to the changes in cellular zinc levels (**Figure 5**). Although the IL2 loop is intrinsically disordered,^27^ the cysteine accessibility experiments, which is sensitive to the local structure and dynamics, allowed us to conclude that the IL2 loop becomes more exposed and/or more disordered as zinc levels increase, providing a reasonable explanation for the zinc dependence of ZIP4 endocytosis (**Figure 6**). According to the proposed elevator transport mode,^54-56^ zinc binding to the transporter triggers a global conformational change from the outward-facing state to the inward-facing state. This is achieved by vertical lifting of the transport domain, a 4-transmembran helix (TM) bundle (TM1/4/5/6, blue domain in **Figure 6**), relative to the scaffold domain (TM2/3/7/8, green domain in **Figure 6**). Since the IL2 loop connects TM3 (scaffold domain) and TM4 (transport domain), the large relative movement between the two domains is likely to be transmitted to the IL2 loop, making the LxL motif more accessible to AP2 as zinc levels rise, thus increasing the chance of ZIP4 being recruited by AP2 for clathrin-mediated endocytosis.

Taken together, we conclude that the LxL motif in ZIP4, the conformation of which responds to changes in cellular zinc levels, functions as an atypical dileucine motif to interact with AP2 in a zinc-responsive manner for clathrin-mediated endocytosis. This work advances the understanding of the molecular basis of the post-translational regulation of ZIP4, an essential mammalian zinc transceptor. The proposed binding mode of the LxL motif to AP2 suggests that the search for sorting signals has not yet completed and that additional sorting signals and thus endocytosis mechanisms may be discovered in other nutrient transceptors that undergo substrate-induced endocytosis.

## EXPERIMENTAL MODEL AND SUBJECT DETAILS

### Cell Lines

Human embryonic kidney cells (HEK293T, ATCC, Cat#CRL-3216) and human hepatoblastoma cell line Hep G2 (HepG2, ATCC, Car#HB-8065, a generous gift from Dr. Hua Xiao, Michigan State University) were cultured in Dulbecco’s modified eagle medium (DMEM, Thermo Fisher Scientific, Invitrogen, Cat#11965092) supplemented with 10% (v/v) fetal bovine serum (Thermo Fisher Scientific, Invitrogen, Cat#15240062) at 5% CO_2_ and 37°C.

### Genes and Pasmids

The complementary DNA of human ZIP4 (GenBank ID: BC062625) from Mammalian Gene Collection was purchased from GE Healthcare. The DNA encoding ZIP4 were inserted in to a modified pEGFP-N1 vector (Clone, Cat#6085-1) in which the downstream EGFP gene was deleted and a human influenza hemagglutinin (HA) tag was inserted before the stop codon, leading to a ZIP4-HA construct. The pCMV3-based vectors for the expression of N-terminal FLAG-tagged mouse ZIP4 (GenBank ID: 72027, Cat#MG53693-NF) and the untagged human AP2S1 (σ2 subunit of AP2, GenBank ID: 1175, Cat#HG12478-UT) in mammalian cells were purchased from Sino Biological. The DNA sequence encoding the ECD of human ZIP4 was amplified and inserted into the pLW01 vector for the expression of N-terminal His_6_-tagged ZIP4-ECD in *E*.*coli*. All the mutations were made using PfuTurbo DNA polymerase (Agulent, Cat#600250) and verified by DNA sequencing. All the plasmids were purified using Miniprep (Promega, cat#A1460) or Maxiprep (QIAGEN, Cat# 12163). All the primer sequences for mutagenesis are listed in **Table S1**.

## METHOD DETAILS

### Transient transfection and RNA interference

For transient expression of ZIP4 and its variants, HEK293T cells were transfected with the corresponding plasmids using Lipofectamine 2000 (Thermo Fisher Scientific, Invitrogen, Cat#11668019) and cultured for 16 h prior to the antibody uptake assay. To enhance knockdown efficiency, HepG2 cells were subjected to sequential siRNA transfection: after the initial transfection, a second round was performed 48 h later, followed by an additional 48 h incubation before analysis. In the σ2 subunit knockdown-rescue experiments, siRNA-resistant σ2 or its variants were co-transfected with σ2-targeting siRNA during the second transfection, and cells were cultured for 48 h before the assay. All siRNA sequences are provided in **Table S1**.

### Western blot

Proteins separated by SDS-PAGE were transferred to PVDF membrane (Millipore, Cat# PVH00010) and blocked with 5% (w/v) non-fat dry milk. The primary antibodies against AP2A1 (α subunit of AP2, MyBioSource, Cat# MBS416635), AP2S1 (Abcam, Cat# ab128950), Clathrin heavy chain (BD Biosciences, Cat# 610499) and ZIP4 (Proteintech, Catalog# 20625-1-AP) were applied at 1:1000 dilution, and the primary antibodies against FLAG tag (Agilent, Cat# 200474) and HA tag (Thermo Fisher Scientific, Cat# 26183) were applied at 1:3000 dilution. As loading control, β-actin levels were detected using an anti-β-actin antibody at 1:2000 (Cell Signalling Technology, Cat# 4970). Primary antibodies were detected with HRP-conjugated secondary antibodies (Anti-rat IgG, Cell Signaling Technology, Cat# 7077S, applied at 1:6000 dilution; Anti-rabbit IgG, Cell Signaling Technology, Cat# 7074S, applied at 1:2500 dilution; Anti-mouse IgG, Cell Signaling Technology, Cat# 7076S, applied at 1:6000 dilution) and signals were generated by chemiluminescence (VWR, Cat# RPN2232). Images were taken using a Bio-rad ChemiDocTM Imaging System.

### Antibody uptake assay

To study the role of the LQL segment in ZIP4 endocytosis, ZIP4 or its variants were transiently expressed in HEK293T cells. Cells were firstly washed by Dulbecco’s phosphate buffered saline (DPBS) (Sigma-Aldrich, Cat#D8537-500ML) once and then trypsinized by 0.25 % Trypsin-EDTA (Gibco, Cat#25-200-056) under 37°C for 2 min. The cells were then suspended in DMEM medium supplemented with 10% FBS, and centrifuged at 40 *g* for 3 min. The cells were then resuspended in an ice-cold 500 µl DMEM medium supplemented with 10% FBS before Alexa Fluor-647 conjugated anti-HA antibody (Invitrogen, Cat#26183-A647; RRID: AB_2610626) was applied at 1:400 dilution. Cells were equally divided into two samples – cells in control group was maintained at 4°C for 45 min, whereas cells in the experimental group were incubated at 37°C for the same period of time. After incubation, cells were centrifuged at 40 *g* and 4°C for 2 min and washed three time by DPBS. Cells were then washed with an acidic buffer (DMEM+0.2% BSA, pH 4.0 adjusted by HCl) for 45 s to strip the surface bound anti-HA antibodies, which was terminated by the addition of ten times the volume of normal DMEM medium. Cells were then washed with DPBS once, fixed with 4% formaldehyde for 15 min at room temperature, and washed three more times with DPBS before flow cytometry analysis.

To study endogenous ZIP4 endocytosis in HepG2 cells, cell suspension was generated by the same approach except for a longer trypsin digestion for 5 min under 37°C. Cells were then incubated with the 1G3B2 anti-ZIP4 mAb at 4 µg/ml under 4°C and 37°C respectively, for 90 min. After washing for three time by DPBS, the surface bound antibodies were stripped with the acidic buffer, followed by fixation and wash as mentioned above. To permeabilize cells and detect the internalized anti-ZIP4 mAb, cells were incubated with Alexa Fluor-568 conjugated goat anti-mouse IgG (Thermo Fisher Scientific, Cat#A-11004) at 1:500 dilution and 0.1% Triton X-100 (americanBIO, Cat#9002-93-1) in DPBS at 4°C for 1 h. Cells were then washed for three times and suspended in DPBS for flow cytometry analysis.

In order to exclude the dead cells in the antibody uptake assay, for both HEK293T cells and HepG2 cells, LIVE/DEAD™ fixable violet dead cell stain (Invitrogen, Cat#L34963) was added to cell suspensions at 1 µl/ml for an incubation with antibodies at 4°C or 37°C.

### Flow cytometry

Cell samples were applied to a ThermoFisher Attune Cytpix flow cytometer. At least 10,000 cells were assessed for each sample. The cellular debris was excluded based on FSC-A vs. SSC-A panel (**Figure S1B**) and single cells were selected through FSC-A vs FSC-H gating. To generate live/dead control group, the same amount of HEK293T or HepG2 cells were placed at either 65°C or 37°C for 15 min, and then mixed and stained by LIVE/DEAD™ stain at 4°C for 30 min. The cells were then washed twice by DPBS before flow cytometry analysis. Based on the result of this live/dead group, the same gating was applied to all the experimental groups. MFIs of Alexa Fluor-647 (HEK293T cells) or Alexa Fluor-568 (HepG2 cells) were calculated for all live and single cells in experimental groups (incubated at 37°C) and control groups (incubated at 4°C). The difference of MFI (ΔMFI) between experimental groups and control groups reports antibody uptake, which serves as an indicator of ZIP4 endocytosis. All the flow cytometry data processing was performed in FlowJo 10.8.1.

### ZIP4 endocytosis detection by immunofluorescence microscopy

To visualize human ZIP4 endocytosis in HEK293T and HepG2 cells, cell suspensions in DMEM medium were treated with either Alexa Fluor-488 conjugated anti-HA antibody at 1:400 (HEK293T) or the 1G3B2 anti-ZIP4 mAb at 4 µg/ml (HepG2). Cell suspensions were incubated at 4°C and 37°C, respectively, for 45 min for HEK293T cells and 90 min for HepG2 cells. After centrifugation, cells were fixed with 4% formaldehyde and washed three times by DPBS. HepG2 cells were further treated with Alexa Fluor-568 conjugated goat anti-mouse IgG at 1: 500 dilution in DPBS plus 0.1% Triton X-100 at room temperature for 1 h. HepG2 cells were then washed for 3 times by DPBS before imaging. The images for HEK293T cells and HepG2 cells are shown in **Figure S1A** and **Figure 2A**, respectively.

To visualize the endocytosis of N-FLAG-tagged mouse ZIP4 (wild type and the CysFree-TMD variant), HEK293T cells attached on poly-D-lysine coated coverslips and transiently expressing mouse ZIP4 were incubated with an anti-FLAG antibody (Agilent, Cat#200474) at 1: 3000 dilution in DMEM medium supplemented with 10% FBS under 37°C for 45 min. Cells were then washed for three times with DPBS and fixed by 4% formaldehyde, followed by permeabilization with 01.% Triton X-100 and incubation with Alexa Fluor-488 conjugated goat anti-rat IgG (Invitrogen, Cat#A-11006) at 1: 200 dilution in DPBS under room temperature for 1 h. After three times of wash with DPBS, the coverslips were mounted on the slides with fluoroshield mounting medium containing DAPI (Abcam, Cat#ab104139). The images are shown in **Figure S4C**.

All images were taken with a 20 x objective using Nikon C2 confocal microscope.

### Cysteine accessibility assay

DMEM medium supplemented with 10% FBS was treated by Chelex® 100 resin (Sigma-Aldrich, Cat#C7901-25G) to generate a Chelex-treated medium. HEK293T cells expressing CysFree-TMD and its variants were firstly washed with the Chelex-treated medium and then incubated with 20 µM N,N,N, N -tetrakis(2-pyridylmethyl) ethylenediamine (TPEN) (Sigma-Aldrich, Cat#P4413) and 0.5% DMSO at 37°C for 15 min. Cells were then washed twice with the Clelex-treated medium for analysis under zinc depletion conditions. To create a zinc repletion condition, after depletion of both extracellular and intracellular zinc pools as mentioned above, 10 µM zinc chloride and 10 µM 2-mercaptopyridine N-oxide sodium salt (pyrithione, Sigma-Aldrich, Cat#H3261-1G) were added to the HEK293T cells. The DMEM medium supplemented with 10% FBS without treatment was used for basal condition.

The treated HEK293T cells were incubated with indicated concentrations of N-ethylmaleimide (NEM, Thermo Scientific, Cat#23030) under 4°C for 1 h. Cells were then washed twice to remove the excessive NEM with a buffer containing 100 mM Tris-HCl, 60 mM NaCl, 10 mM KCl, pH 7.0, and lysed with 6 M urea, 0.5% SDS, and 0.5 mM DTT (to quench residual NEM). Cell lysates were then heated for 10 min under 96°C to completely denature proteins, followed by an incubation with 5 mM monofunctional PEG maleimide 5k (mPEG5k, Creative PEGWorks, Cat#PLS-234) to react with free thiol cysteine residue for 1Uh at room temperature with gentle shaking. Samples were then mixed with SDS sample loading buffer containing 20% β mercaptoethonal (β-ME) and subjected to SDS-PAGE. CysFree-TMD variants were detected in Western blot using the anti-FLAG antibody.

### Zinc transport assay and inductively coupled plasma mass spectrometry (ICP-MS)

^70^Zn transport assay and ICP-MS analysis were conducted as described.^57^ In brief, 16 hours post transfection, HEK293T cells expressing mouse ZIP4 (wild type and the CysFree-TMD variant) were washed with a washing buffer (10 mM HEPES, 142 mM NaCl, 5 mM KCl, 10 mM glucose, pH 7.3), followed by incubation with the Chelex treated medium. 5 µM ^70^Zn was added to cells to initiate zinc transport. After incubation at 37°C for 30 min, cells were transferred on ice and an equal volume of the ice-cold washing buffer containing 1 mM EDTA was added to the cells to terminate metal uptake. After three washes with the ice-cold washing buffer, 70% trace nitric acid was added to cells for digestion and subsequent ICP-MS analysis. Zinc uptake by cells was expressed as the count ratio of ^70^Zn/^31^P determined by ICP-MS, and zinc transport activity was calculated by subtracting zinc uptake by the cells transfected with an empty vector from that by the cells expressing metal transporters.

### Production of anti-human ZIP4 monoclonal antibodies

His_6_-tagged human ZIP4-ECD was expressed and purified in the same way as bat ZIP4-ECD.^45,46^ In brief, *E*.*coli* strain of Origami B(DE3) pLysS (Novagen, Cat#70839) was cultured in lysogeny broth medium and protein expression was induced by 0.2 mM isopropyl-beta-D-thiogalactopyranoside, followed by culturing at 16°C overnight. ZIP4-ECD in cell lysate was purified using nickel-nitrilotriacetic acid affinity column and applied to size-exclusion chromatography after the His_6_-tag was removed by thrombine. The purified protein was delivered to Creative Biology to immunize mice. Hybridomas were screened and positive clones, including Clone 1G3B2, were identified and verified using enzyme-linked immunosorbent assay.

Hybridoma (Clone 1G3B2) were cultured in Hybridoma-SFM medium (Thermo Fisher Scientific, Cat#12045076) and the culture supernatant was collected and clarified by centrifugation at 6000 rpm at 4°C for 10 min, followed by filtration through a 0.22 μm membrane. The clarified supernatant was loaded onto a Protein G Agarose (Thermo Scientific, Cat#20398) pre-equilibrated with Protein G IgG Binding Buffer (Thermo Scientific, Cat#21011). After washing with ten column volumes of IgG Binding Buffer to remove non-specifically bound proteins, bound IgG was eluted with 0.1 M glycine-HCl buffer (pH 2.9) and immediately neutralized with 1 M Tris-HCl (pH 8.0). Eluted fractions were pooled and concentrated using Amicon Ultra centrifugal filters (Millipore, 30 kDa MWCO), and applied to buffer exchange into phosphate-buffered saline. The final IgG concentration was determined by the absorbance at 280 nm measured using a NanoDrop spectrophotometer. Purity of 1G3B2 mAb and its specific staining of native human ZIP4 expressed in HEK293T cells are shown in **Figures S2A & S2B**, respectively.

## QUANTIFICATION AND STATISTICAL ANALYSIS

We assumed a normal distribution of the samples and multiple comparisons were assessed using two-tailed Student’s *t-*test. A *P*< 0.05 was considered statistically significant. Data were presented as mean ± standard deviation.

## RESOURCE AVAILABILITY

Lead Contact and Materials Availability

Further information and requests for resources and reagents, including the plasmids and the 1G3B2 anti-ZIP4 antibody generated in this study, should be directed to the Lead Contact, Dr. Jian Hu and will be fulfilled upon completion of a Materials Transfer Agreement.

### Data Availability

Any additional information required to reanalyze the data reported in this paper is available from the lead contact upon request.

## ACKNOWLEDGEMENT

We thank Flow Cytometry Core Facility at Michigan State University in training and assistance of data collection and analysis. We thank Quantitative Bio Element Analysis and Mapping (QBEAM) center at Michigan State University for the assistance of using ICP-MS. We thank Dr. Yuhan Jiang at Michigan State University in measuring the zinc transport activity of mouse ZIP4. This work was supported by NIH grant GM140931 (to J.H.).

## AUTHOR CONTRIBUTIONS

Investigation: T.W. and C.Z. conducted cell biological experiments and data analysis, T.W. conducted biochemical experiments and data analysis, Y.Z. generated the 1G3B2 mAb. Writing: J.H., T.W., C.Z., and Y.Z.; Conceptualization: J.H., T.W., and C.Z.; Supervision: J.H.

