## Supplementary Information for "Endocytosis of a zinc transceptor ZIP4 is mediated by AP2 through an atypical dileucine motif"

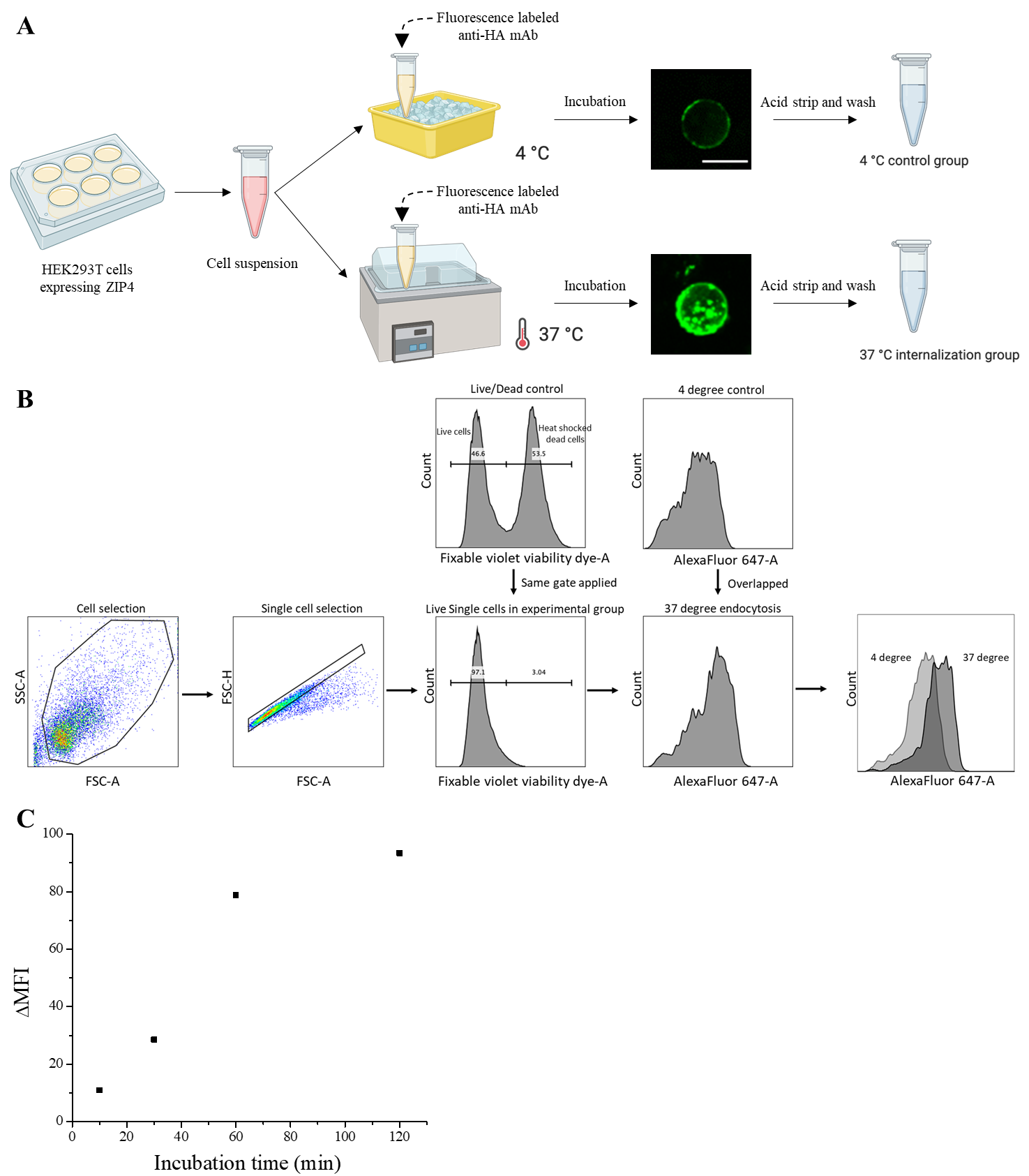

**Figure S1.** Flow cytometry-based human ZIP4-HA endocytosis analysis. (**A**) Cartoon illustration of the antibody uptake assay. The confocal images of HEK293T cells that express ZIP4-HA after incubation with Alexa Fluor 488-labeled anti-HA antibodies at 4°C and 37°C, respectively, are shown. The scale bar is 25 µm. (**B**) Flow chart for flow cytometry data processing. (**C**) Time course of Alexa Fluor 647-labeled anti-HA antibody uptake by HEK293T cells expressing ZIP4-HA. δMFI was calculated by subtracting the MFI of the cells incubated with the antibody at 4°C from the MFI of the cells incubated with the antibody at 37°C.

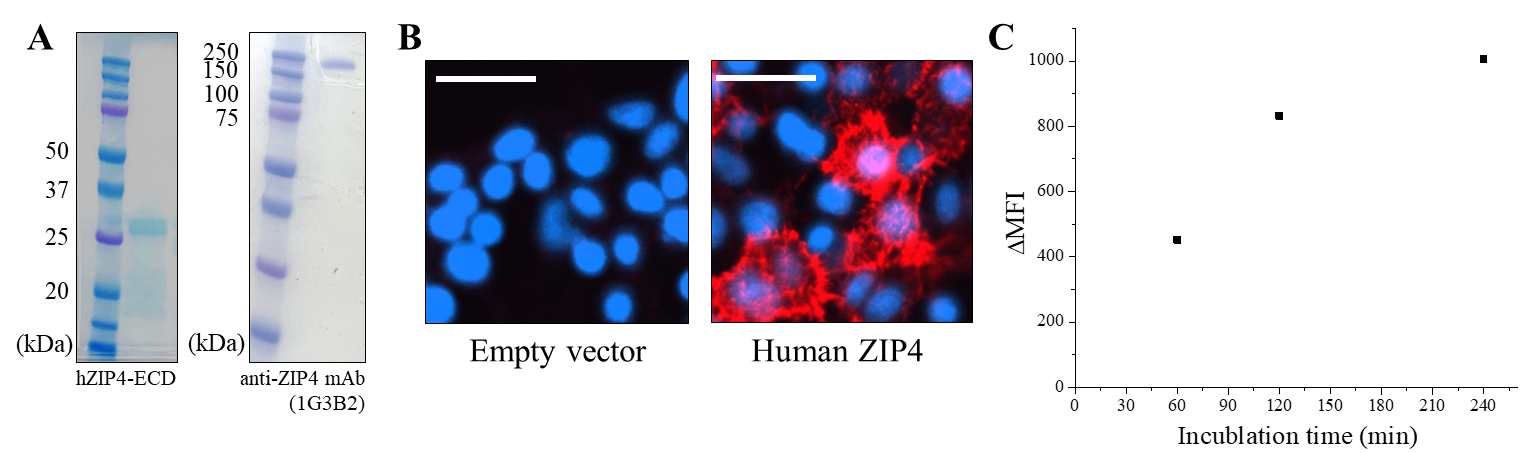

**Figure S2.** Generation of the 1G3B2 anti-human ZIP4 mAb to study ZIP4 endocytosis. (**A**) SDS-PAGEs of purified hZIP4-ECD used as antigen for mice immunization (*left*) and purified 1G3B2 anti-hZIP4 mAb from hybridomas. (**B)** Immunofluorescence imaging of HEK293T transfected with an empty vector (*left*) or a vector harboring ZIP4-HA (*right*). 1G3B2 was used as the primary antibody and an Alexa Fluor 568-conjugated anti-mouse antibody was used the secondary antibody. The scale bar represents 25 µm. (**C**) Time course of 1G3B2 uptake by HepG2 cells. δMFI was calculated by subtracting the MFI of the cells incubated with the antibody at 4°C from the MFI of the cells incubated with the antibody at 37°C.

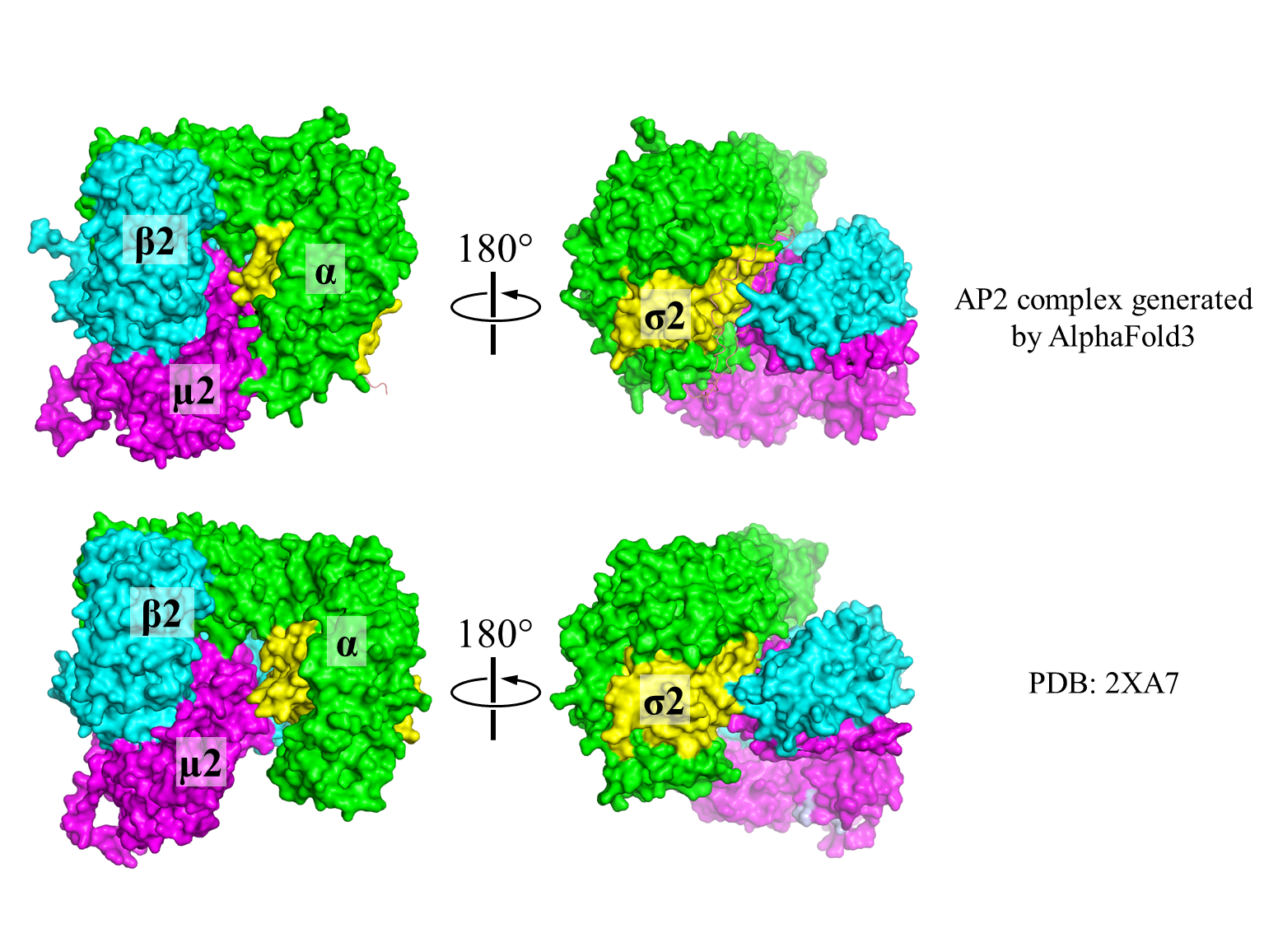

**Figure S3.** Structural comparison of the structural model of AP2 predicted by AlphaFold 3 with the experimentally solved AP2 structure in the open state (PDB: 2XA7).

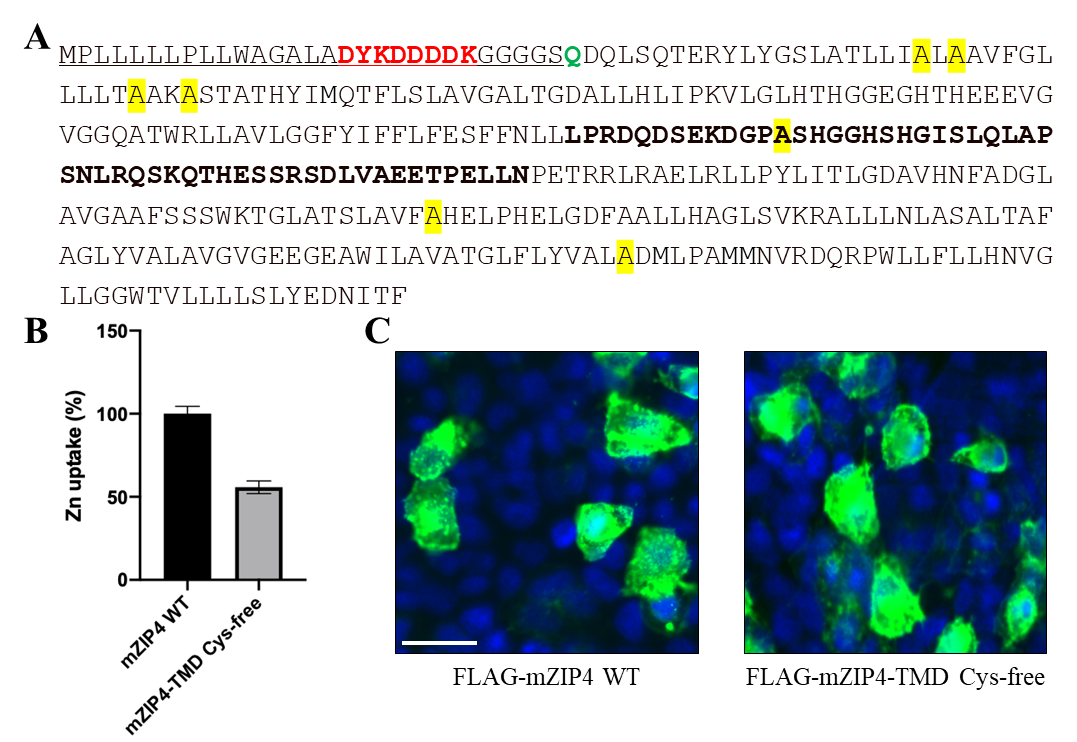

**Figure S4.** Characterization of the mouse ZIP4-TMD Cys-free variant (CysFree-TMD). (**A**) Amino acid sequence of the CysFree-TMD with an N-terminal FLAG tag (red). The alanine residues introduced to replace cysteine residues are highlighted in yellow. The amino acid sequence of IL2 is bolded. The underlined sequence is from the vector purchased from Sino Biological Inc. and contains a universal signal peptide for membrane protein expression. The first residue from mouse ZIP4 is a glutamine (green). (**B**) Cell-based transport assay of wild-type mZIP4 and the CysFree-TMD variant. Activity of the latter is expressed as a percentage of the wild-type transporter. The error bar indicates 1±S.D. (n=3). (**C**) Immunofluorescence imaging of FLAG-mZIP4 and the FLAG-CysFree-TMD variant transiently expressed in HEK293T cells. Live cells were incubated with an anti-FLAG antibody at 37°C for 30 min to allow antibody internalization, and then fixed and permeabilized for the staining with an AlexaFluor-488 conjugated anti-rat antibody. The scale bar indicates 20 μm.

**Table S1.** Primers and siRNA nucleotides used in this study.

| Primer | Sequence (forward only, 5’-3’) |
| --- | --- |
| **Human ZIP4-HA** |  |
| L452A | GGCCACAGCCACGGTGTGTCCGCCCAGCTGGCACCCAGCGAGCTC |
| L452N | GGCCACAGCCACGGTGTGTCCAACCAGCTGGCACCCAGCGAGCT |
| L452I | GGCCACAGCCACGGTGTGTCCATCCAGCTGGCACCCAGCGAGCTC |
| L452M | GGCCACAGCCACGGTGTGTCCATGCAGCTGGCACCCAGCGAGCTC |
| Q453A | CACAGCCACGGTGTGTCCCTGGCCCTGGCACCCAGCGAGCTCCGG |
| Q453N | CACAGCCACGGTGTGTCCCTGAACCTGGGCACCCAGCGAGCTCCGG |
| Q453E | CACAGCCACGGTGTGTCCCTGGAGCTGGGCACCCAGCGAGCTCCGG |
| Q453K | CACAGCCACGGTGTGTCCCTGAAGCTGGGCACCCAGCGAGCTCCGG |
| Q453L | CACAGCCACGGTGTGTCCCTGCTGGGCACCCAGCGAGCTCCGG |
| L454A | AGCCACGGTGTGTCCCTGCAGGCCGCACCCAGCGAGCTCCGGCAG |
| L454N | AGCCACGGTGTGTCCCTGCAGAACGCACCCAGCGAGCTCCGGCAG |
| L454I | AGCCACGGTGTGTCCCTGCAGATCGCACCCAGCGAGCTCCGGCAG |
| L454M | AGCCACGGTGTGTCCCTGCAGATGGCACCCAGCGAGCTCCGGCAG |
| **AP2-σ2** |  |
| siRNA-resistant | GCCGTGGTCACCGTCCGCGATGCTAAGCATACCAACTTTGTGGAGTTC |
| V88D | GCCATTCACAACTTCGATGAGGTCTTAAACGAA |
| L103S | GTCTGTGAACTGGACAGCGTGTTCAACTTCTAC |
| N9S | TTTATCCTCATCCAGAGCCGGGCAGGCAAGACG |
| R15S | CGGGCAGGCAAGACGAGCCTGGCCAAGTGGTAC |
| E100S | TTCCACAATGTCTGTAGCCTGGACCTGGTGTTC |
| **Mouse ZIP4** |  |
| TMD (Δ1-327) | GATGACGACGATAAGGGTGGAGGCGGTAGCCAGGACCAGCTCAGTCAAACAGAGAGGTAT |
| C348A/C350A | GCCACCCTGCTCATCGCCCTCGCTGCTGTGTTCGGTCTT |
| C360A/C363A | CTTCTGCTGCTGACCGCTGCCAAAGCCAGCACAGCCACCCAC |
| C454A | GAGAAAGATGGGCCTGCTAGCCATGGTGGGCAC |
| C548A | TCATTGGCGGTGTTCGCTCATGAGCTGCCCCAT |
| C616A | CTTTACGTGGCGCTTGCTGACATGCTCCCAGCC |
| CysFree-TMD R444C | AACCTCTTGTTGCCCTGCGACCAGGATTCTGAG |
| CysFree-TMD E449C | AGCAGGCCCATCTTTGCAAGAATCCTGGTCCCT |
| CysFree-TMD A454C | GAGAAAGATGGGCCTTGCAGCCATGGTGGGCAC |
| CysFree-TMD I463C | GGGCACAGCCATGGATGCTCTCTGCAGCTGGCA |
| CysFree-TMD N471C | CAGCTGGCACCAAGCTGCCTCCGACAGTCCAAA |
| CysFree-TMD T478C | CGACAGTCCAAACAGTGCCATGAAAGCTCTCGT |
| CysFree-TMD A488C | CGTTCAGACTTGGTGTGCGAGGAGACCCCGGAA |
| CysFree-TMD N496C | ACCCCGGAACTACTGTGCCCAGAGACCCGGCGA |
| **siRNA nucleotides** |  |
| AP2-α (GeneGlobe ID: SI04146429) | cgcggtgttcatctccgacat |
| AP2-σ2  (GeneGlobe ID: SI00297472) | ccgagacgccaaacacaccaa |
| Clathrin heavy chain (GeneGlobe ID: SI00299880) | taatccaattcgaagaccaat |
| Human ZIP4 (GeneGlobe ID: SI00724598) | caggaccagctcagccagtca |
| Control siRNA (Qiagen, Cat# 1022076) | aattctccgaacgtgtcacgt |
